# The microbiome of *Lotus* nodules varies with plant health in a species-specific manner

**DOI:** 10.1101/2021.05.19.441130

**Authors:** Duncan B. Crosbie, Maryam Mahmoudi, Viviane Radl, Andreas Brachmann, Michael Schloter, Eric Kemen, Macarena Marín

## Abstract

Nitrogen fixation is carried out inside nodules of legumes by symbiotic rhizobia. Rhizobia dominate the nodule microbiome, however other non-rhizobial bacteria also colonise root nodules. It is not clear whether these less abundant nodule colonisers impact nodule function. In order to investigate the relationship between the nodule microbiome and nodule function as influenced by the soil microbiome, we used a metabarcoding approach to characterise the communities inside *Lotus burttii, Lotus japonicus* and *Lotus corniculatus* nodules from plants that were either starved or healthy resulting from inoculations with different soil suspensions in a closed pot experiment. We found that the nodule microbiome of all tested *Lotus* species differed according to inoculum, but only that of *L. burttii* varied with plant health. Using a machine learning algorithm, we also found that among the many non-rhizobial bacteria inside the nodule, amplicon sequence variants that were related to *Pseudomonas* were the most indicative signatures of a healthy plant nodule microbiome. These results support the hypothesis that legume nodule endophytes may play a role in the overall success of root-nodule symbiosis, albeit in a plant host specific manner.

## INTRODUCTION

Leguminous plants have evolved a mutualistic interaction with nitrogen-fixing rhizobia in which the bacteria are hosted and nourished in root organs called nodules in exchange for ammonia. This so-called root-nodule symbiosis is initiated by a two-way signalling between the symbiosis partners, which activates distal cell divisions in the root cortex and culminates in the formation and infection of nodules (1). Here the bacteria differentiate into plant-dependent, nitrogen-fixing endosymbiotic bacteriods (2). The fixation of nitrogen is an energetically expensive process for the host, which required at least 16 ATP molecules per N_2_ molecule to fuel the nitrogenase enzyme produced by the rhizobia (3). Thus, to prevent infection of the carbon-rich nodules by pathogens, host plants have evolved complex recognition mechanisms that ensure symbiotic specificity (4).

Root-nodule symbiosis is highly species specific and many plants will only form an effective symbiosis with a narrow range of rhizobia (5). Even within these pairings there is variation in nitrogen fixation efficiency (6). Some bacteria can also nodulate plants and not fix any nitrogen at all (7). Examples of ineffective nitrogen fixation have been described after the introduction of crop legumes into areas where previously native legumes grew. For instance, inefficient nitrogen fixation occurs in fields where perennial and annual clovers co-exist (7). Native rhizobial species associated with native legumes can outcompete inoculant strains (8). In extreme cases endogenous rhizobia can completely block the nodulation of introduced rhizobia. For example, the nodulation of the pea cultivars Afghanistan and Iran by rhizobial inoculants is suppressed in natural soils by the presence of a non-nodulating strain (9). This suggests that interactions of the soil microbiota with the host plant are critical for the establishment of efficient nodules. However, we are far from understanding what factors determine the success of single microbes to compete for resources at the plant soil interface, in particular nodule endophytes and how these affect nitrogen fixation effectiveness.

There is clear evidence to suggest that the host controls the makeup of the microbiota in its vicinity. *Lotus japonicus* selects for a broad taxonomic range of bacteria, in addition to the symbiont, within the rhizosphere, endosphere, and the nodule. This selectivity filters the diverse soil microbiome into a distinct and taxonomically narrow community within the nodule (10). Despite this selective pressure, non-nodulating bacteria, such as *Pseudomonas* sp., *Klebsiella* sp. and *Rhodococcus* sp. have been isolated from plant nodules (11-13). Although these isolates do not directly nodulate the plant, they contribute to plant growth in a number of ways, such as increasing the availability of soluble phosphate and producing plant compounds beneficial for plant growth like siderophores and indoleacetic acid (14-16). In addition, non-rhizobiales microbes found in nodules of *Medicago truncatula* produce antimicrobial compounds that may shape the community and the overall function of the nodule microbiome (17). Microbe-microbe interactions could also impart an effect on the overall functionality of the symbiosis, for instance mediated by antimicrobial activity (18), suppression of plant pathogens (19), or by horizontal gene transfer (20). Although these complex interactions could dictate the effectiveness and specificity of the symbiosis, little is known about how rhizobia interact with other members of the nodule microbiome.

In this work we determined the nodule microbiome of three *Lotus* species upon inoculation with soil suspensions that led to the growth of either starved or healthy plants. We used metabarcoding-based high-throughput sequencing to characterise the microbiome in nodule samples that varied in plant species origin, soil inocula and plant health. Network analyses and machine learning algorithms identified microbiome members specifically associated to nodules of healthy, but not of starved *L. burttii* plants. Our results suggest that although root nodule symbiosis is a binary interaction, there are other microbes which modulate this mutualistic interaction.

## MATERIALS AND METHODS

### Soil collection and inoculum preparation

Soil samples were collected from two neighbouring sites in a semi-urban area south west of Munich, Germany. Site 1 (48 06’29.9”N, 11 27’38.9”E) has consistently been home to wild *Lotus corniculatus* whereas site 2 (48 06’33.2”N, 11 27’41.4”E) has been subjected to tilling and physical disturbance and did not contain *Lotus* plants at the time of collection. Soil samples were taken from the top layer (0-20 cm deep) after plant material was removed from the site in May 2019 and October 2018. Physicochemical property measurements of each soil were performed by AGROLAB Agrarzentrum GmbH (Landshut, Germany). Soil samples were sieved to remove stones and plant material with a 2 mm sieve, mixed 1:5 with a nitrogen-limiting FAB liquid medium, and stirred for 2 h. Soil particulate matter was removed by centrifugation at 1 000 g for 5 min. Soil suspensions were used as inputs and a qPCR was run to compare the quantity of soil bacteria present in both soil suspensions.

### Plant growth and inoculation conditions

*Lotus burttii* B-303 (seed bag no. 91105), *Lotus japonicus* Gifu B-129 (seed bag no. 110913) and *Lotus corniculatus* cv. Leo (Andreae Saaten, Regensburg, Germany) seeds were scarified and then sterilised by incubation in a sterilising solution (1.2 % NaOCl, 1 % SDS) for 8 min before being washed 3 times with sterile water. Seeds were then soaked in sterile water for 2-3 h and germinated on 0.5 B5 agar medium (21) for 3 days in dark followed by 3 days in a long-day photoperiod (16 h light, 8 h dark) at 24°C. Seedlings were then transferred into sterilised tulip-shaped Weck jars (10 seedlings per jar) containing 300 mL of a sand:vermiculite mix (1:2) and supplemented with 40 ml of a low nitrogen FAB medium, to create nitrogen-limiting growth conditions as described before (22). Jars were sealed with micropore tape to create a closed system. Seedlings were left to recover for 2 days in a long-day photoperiod. After the 2-day recovery, each seedling was inoculated with 1 mL of soil suspension. *L. burttii* and *L. japonicus* treatments consisted of 150 plants from 3 independent experiments, and *L. corniculatus* treatments consisted of 50 plants per condition from 1 independent experiment.

### Harvesting, phenotyping, and nodule surface sterilisation

Plants were harvested and phenotyped 5 weeks post inoculation. Shoot length, shoot dry weight, nodule number, nodule colour and plant health were recorded. Roots were removed from shoots and sonicated using the Bioruptor® (Diagenode, Seraing, Belgium) twice for 15 min. Nodules from 3-4 plants were excised and pooled based on similarity of plant shoot and nodule phenotype. Pooled nodules were treated with 70 % ethanol for 1 min followed by 2 % NaOCl for 2.5 min. Nodules were then washed with sterile water 8 times and after the removal of the final water wash, samples were snap frozen in liquid N_2_. The final wash was plated onto 20Q supplemented with 3.8 % w/v Mannitol (modified from (23)) agar to assess sterilisation.

### DNA extraction

Nodule samples were homogenised six times in a Mixer Mill 400 (Retsch, Haan, Germany) at a frequency of 30 s^-1^ for 1 min. DNA was then extracted according to a modified protocol from Töwe et al., 2011 (24). For extraction of DNA from the inputs, soil suspensions were centrifuged at 5 000g and DNA from pellets was extracted according to the CTAB method described by the Doe Joint Genomics Institute (25). The concentrations of extracted DNA samples were quantified with a Qubit 2.0® fluorometer (Invitrogen, Carlsbad, California, United States).

### Quantitative PCR

Quantitative PCR (qPCR) was performed using the FP *16S rDNA* (5’-GGTAGTCYAYGCMSTAAACG-3’) and RP *16S rDNA* (5’-GACARCCATGCASCACCTG-3’) primers (26). The 25 µL PCR mixture contained SYBR Green, 2 µL template DNA, 7.5 µL Milli-Q water, 1 µL 10 µM of primers and 1 µL 15 % Bovine Serum Albumin (BSA). The mixture was amplified using a CFX96 Real-Time System (Bio-Rad, Hercules, California, United States) under the following conditions: template was denatured at 94°C for 10 min before 40 cycles of 95°C for 20 s, 57°C for 30 s and 72°C for 45 s, followed by dissociation curve steps of 95°C for 15 s, 60°C for 30 s and 95°C for 15 s. Quantification of the 16S rRNA gene molecules were correlated with a calibration curve constructed with known amounts of a 16S rRNA gene standard plasmid constructed of a *Mesorhizobium septentrionale* 16S rRNA gene sequence cloned into a pUC57 plasmid.

### Amplification, library preparation, and sequencing

To determine bacterial diversity, a metabarcoding approach was utilised. The hypervariable region V3-V4 of the 16S rRNA gene was amplified using universal bacterial primers 335F (5’-CADACTCCTACGGGAGGC-3’) and 769r (5’-ATCCTGTTTGMTMCCCVCRC-3’) fused to Illumina adapters. The primers are specific for bacterial DNA and do not amplify plastidial and mitochondrial plant DNA (27). Amplification reaction volumes were 25 μL using 1 unit of Phusion polymerase, 5 μL 5x High-Fidelity Phusion buffer, 7.5 μL of 1 % BSA, 0.5 μL of 10 mM dNTPs, 0.5 μL of 50 mM MgCl_2_, 0.5 μL of 10 pmol/μL primer and 5 ng of template DNA. The assay was conducted in triplicate under the following conditions: template was denatured at 98°C for 1 min, then 25 cycles of 98°C for 10 s, 55°C for 30 s and 72°C for 30 s, followed by a final step at 72°C for 5 min. PCR products were verified via gel electrophoresis, pooled, and cleaned using CleanPCR beads (CleanNA, Waddinxveen, The Netherlands). Fragments were then indexed with 10 nucleotide barcode sequences using Nextera XT Index Kit v2 Set D primers (Illumina, San Diego, California, USA). Indexing PCR reactions were run in triplicate with a volume of 25 μL using 12.5 μL NEB Next High-Fidelity Master Mix, 2.5 μL of each delegated primer and 20 ng of amplicon under the following conditions: template was denatured at 98°C for 30 s, then 8 cycles of 98°C for 10 s, 55°C for 30 s and 72°C for 30 s, followed by a final step at 72°C for 5 min. PCR products were pooled and cleaned with CleanPCR beads (CleanNA). Quantification and quality control were conducted using AATI Fragment Analyzer (Santa Clara, California, United States). All samples were pooled at an equimolar concentration for paired end 2 × 300 bp sequencing via the MiSeq system (Illumina) using the MiSeq Reagent Kit v3 (600 cycles), as per the manufacturer’s recommendation.

### Sequence and statistical analysis

An average of approximately 141 000 raw Illumina reads per sample were obtained, which were then demultiplexed and had adapter and barcode sequences removed using Cutadapt v

3.1 (28). Reads were then trimmed, merged and filtered using DADA2 (29) in R. The criteria for filtering were minimum lengths of 280 bp for the forward reads and 160 bp for the reverse, as these lengths corresponded to a minimum quality score of 25. Merged sequences were removed of chimeras and chloroplastic and mitochondrial sequences. Amplicon sequence variants (ASVs) were assigned in R using the Silva database (30).

The *phyloseq* V 1.26.1 package in R (31) pipeline was used to infer alpha diversity of ASVs rarefied corresponding to the sample with the lowest number of reads. Multidimensional Scaling using Bray-Curtis (32) distance was performed using *phyloseq* V 1.26.1 package in R (31) in order to assess the beta diversity of microbial communities. Comparisons were visualised using *ggplot* (33) in R and tested for statistical significance (*adonis* test, p<0.01) via Permutational Multivariate Analysis of Variance (PERMANOVA) utilising 999 permutations in the *vegan* package (34). Relative abundance of each genera per sample was calculated using transformed count data. To further specify the composition of the sample microbiome the relative abundance of the most prevalent ASVs (abundance > 0.1%) was calculated for each sample. All abundance levels were calculated using the *phyloseq* V 1.26.1 package (31) in R.

### Machine learning model

ASV tables were filtered to ASVs present in ≥ 50 reads in *L. burttii* plants samples. A support vector machine learning model by svm.SVC(kernel=linear) in python scikit-learn was used to discriminate between starved and healthy plant samples of *L. burttii* on relative abundance filtered ASVs using 5-fold cross validation. svm.coef_function was performed to select for important features.

### Microbial correlation networks

Filtered ASV tables (ASV raw abundances) were used to calculate microbial correlation networks among ASVs using the SparCC (35) algorithm in FastSparc (36). Pseudo *P*-values were inferred from 1 000 bootstraps. Only correlations with *P* < 0.01 were kept for further analyses. Network visualisation was done in Cystoscope v.3.8.2 (37).

## RESULTS

### Species specific effect of soil inoculum on *Lotus* plant growth

Two different soil suspensions were used to inoculate *Lotus burttii, Lotus japonicus*, and *Lotus corniculatus* plants. These *Lotus* species were selected as they all belong to the *L. corniculatus* clade (38), but nodulate with a different range of microsymbionts (39, 40). The first soil (soil 1) was collected at a site that contained healthy wild growing *Lotus corniculatus* plants, while the second soil (soil 2) site contained no leguminous plants at all. A physicochemical soil analysis of the two sites showed minor differences in mineral content as well as in the grain size of the soils (**Table S1**). Soil suspensions were used as inputs and a qPCR was run to compare the quantity of soil bacteria present in the two soil suspensions. Soil suspensions 1 and 2 contained 1.62×10^5^ and 2.28×10^5^ molecules of the 16S rRNA gene per nanogram of extracted DNA, respectively.

Plants were inoculated with each soil suspension and harvested after 5 weeks across 5 independent experiments (**Figure S1**). *L. japonicus, L. burttii* and, to a lesser extent *L. corniculatus*, produced exclusively healthy plants (green leaves, elongated shoots) when inoculated with soil 1 suspension (**Figure 1**). Contrastingly there was marked variation in the shoot growth phenotype seen in all species when inoculated with the soil 2 suspension. Growing alongside the healthy plants was a large contingent of starved plants presenting with shorter shoots and yellow leaves (**Figure 1**). Nodule number also varied dependent on soil suspension inoculum. Plants inoculated with soil 1 consistently developed a higher number of nodules per plant across all species (**Figure S2**). Starved plants inoculated with the soil 2 suspension exhibited two distinct root phenotypes, with and without nodules. The ratio of these phenotypes varied across each plant species. In *L. burttii*, 73.8 % of starved plants contained nodules, while in *L. japonicus* and *L. corniculatus*, 45.2 % and 59.3 % exhibited nodules, respectively (**Figure S3**). However, the most striking difference was that in *L. burttii* 61.3 % of the starved-nodulated plants harboured white nodules, while in the other species most of the few nodules were pink. The pink colouration indicates the presence of leghemoglobin, a pre-requisite for nitrogen fixation (41). Thus, the lack of leghemoglobin in white nodules indicates an environment in which rhizobia cannot fix nitrogen. These results show that the microbiota of the soil 2 suspension is capable of mediating both effective and ineffective symbiosis, although the frequency at which each plant succumbs to an ineffective nodulation differs.

**Figure 1.**
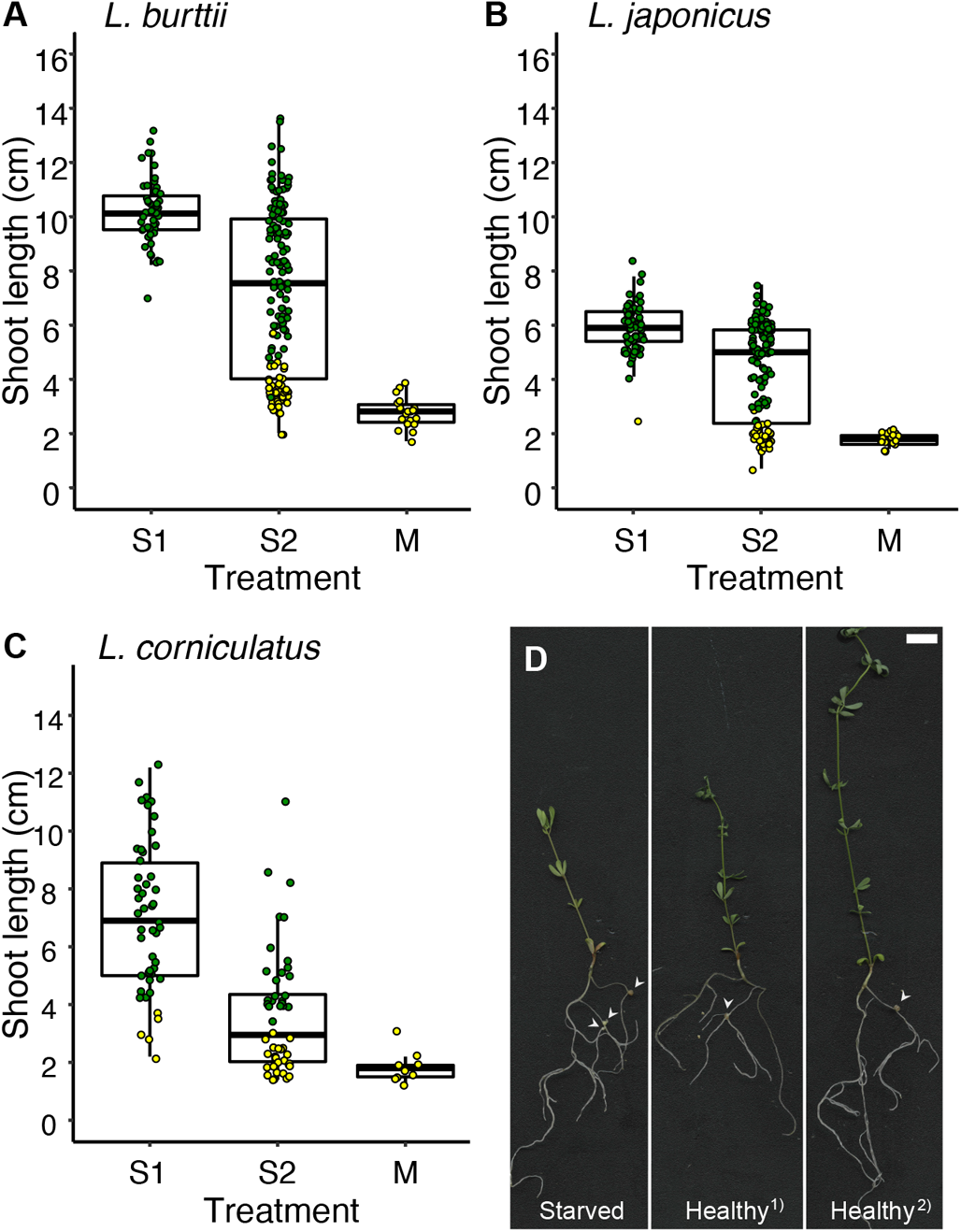
Shoot growth phenotype of *Lotus* plants inoculated with Munich soil suspensions. Shoot growth quantification of *L. burttii* **(A)**, *L. japonicus* **(B)**, and *L. corniculatus* **(C)** plants 5 weeks post inoculation with soil suspensions 1 (S1) and 2 (S2) and a mock (M) treatment. Green and yellow dots indicate plants with healthy and starved phenotypes, respectively. Box plots display the results of 50-150 plants per condition. A total of 49 mock treated plants were included. **(D)** Scanned images of *L. burttii* 5-weeks post inoculation with soil suspension 2. Starved plants exhibited pale green leaves despite having nodules on their roots. The shoots of healthy dark green plants varied in length. Phenotypic variation is depicted in ^1)^ and ^2)^. White arrowheads indicate the position of nodules on plant roots. Plots show the results from one representative experiment. Scale bar: 1 cm.

### Richness and diversity of the *Lotus* nodule microbiome

The microbiome of an effective plant nodule is typically dominated by the respective symbiont, although there can also be minor colonisation by other microbes (13). To investigate how the nodule microbiota varied depending on the plant host, inoculum, and nodule phenotype we sequenced the microbiome of nodules collected from healthy and starved *Lotus* of different species inoculated with different soil suspensions. A variable region of the 16S rRNA gene was sequenced and the output reads were processed, sorted into ASVs, and assigned a taxonomy. ASVs were used as they provide a finer resolution than Operational Taxonomic Units (OTUs) (29). This is important because the 16S rRNA gene of some rhizobia, such as *Mesorhizobium*, can be more than 99% identical between different species (42). In order to capture as much of the diversity that was present, reads were clustered based on 100% similarity. Sequencing produced 13 989 700 paired-end reads after quality filtering, which clustered into 67 442 unique ASVs. Sequence coverage varied between sample types with the nodule samples having an average of 148 679 per sample and the input soil suspension samples having an average of 67 618 reads per sample (**Table S2**). All rarefaction curves reached a saturation plateau (**Figure S4**).

To assess the effect of the host genotype and the inoculum on the nodule microbiome diversity, the alpha and beta diversities of the different nodule samples from all three species were determined. Within sample variation (alpha diversity) was calculated using the Shannon diversity index, which was found to be much higher in the soil input samples compared to the nodule samples (**Figure 2**). The soil suspensions 1 and 2 did not significantly vary in their alpha diversities (Welch two sample t-test, p=0.749) although it was found that plants inoculated with soil suspension 1 produced nodules with a much higher alpha diversity compared to those inoculated with soil suspension 2. This observation was most pronounced in *L. japonicus* and *L. corniculatus* (**Figure 2**). A similar trend in alpha diversity was seen when considering observed ASVs (**Figure 2**).

**Figure 2.**
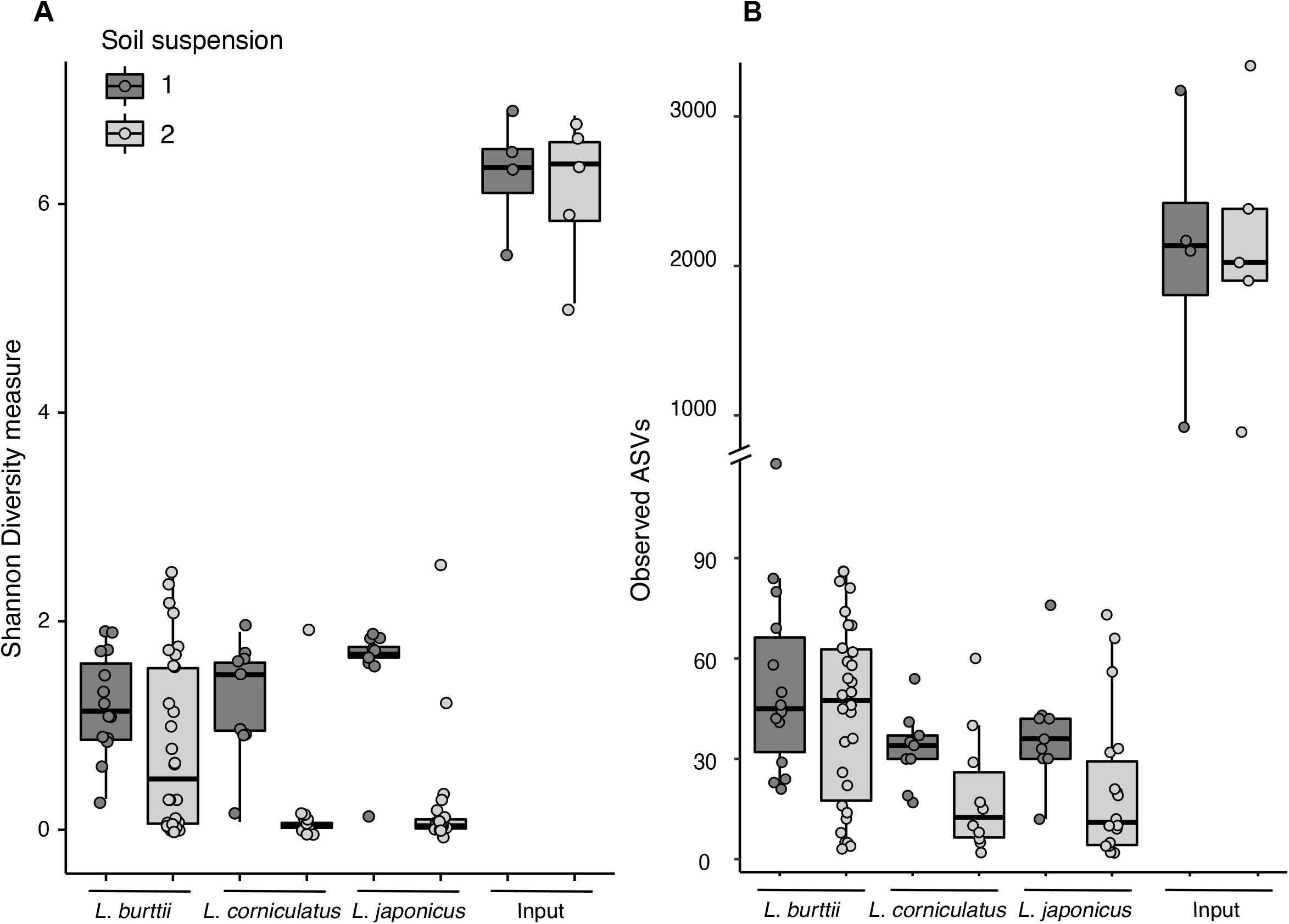
Nodule microbiome alpha diversity plotted by species and soil input. **(A)** Shannon diversity measures and **(B)** observed ASVs of all 99 samples were calculated using unfiltered data.

### Community structure of the nodule microbiome

To analyse the diversity between sample types (beta diversity), principal coordinate and PERMANOVA analyses were conducted based using the Bray-Curtis dissimilarity. A global comparison of the nodule diversity showed an overall separation based on soil input (**Figure S5**; Soil S1 v Soil S2, Pr(>F) = 0.001; **Table S3**). The two soil inputs showed slight differences between one another, although this was not significant (Soil Input S1 v Soil Input S2, Pr(>F) = 0.072, **Table S3**). The most pronounced difference was between nodules of *L. burttii* plants (*Lb* healthy plants - Soil S1 v Soil S2, Pr(>F) = 0.001). In addition, the nodule microbiome makeup of *L. burttii* significantly depended on the plant health status (*Lb* Soil S2 - healthy v starved plants, Pr(>F) = 0.001; **Figure 3, Table S3**), however this was not the case in *L. japonicus* or *L. corniculatus* (*Lj* Soil S2 – healthy v starved plants, Pr(>F) = 0.097; *Lc* Soil S2 – healthy v starved plants, Pr(>F) = 0.742, **Table S3**). The only comparisons between the nodule microbiomes of plants inoculated with both soil suspensions that revealed differences dependent on the host genotype were *L. burttii* and *L. japonicus* inoculated with soil suspension 2 (Soil S2 - *Lb* v *Lj*, Pr(>F) = 0.001, **Table S3**) and *L. japonicus* and *L. corniculatus* (Soil S1 - *Lc* v *Lj*, Pr(>F) = 0.042). As a control, we compared the microbiome of laboratory grown *L. corniculatus* plants to the microbiome of nodules collected from *L. corniculatus* plants growing on site 1 (*Lc* Soil S1 - lab grown v wild plants, Pr(>F) = 0.342) (**Table S3**). These did not significantly differ, supporting that nodules produced in this growth/inoculation system are representative of nodules grown in the wild.

**Figure 3.**
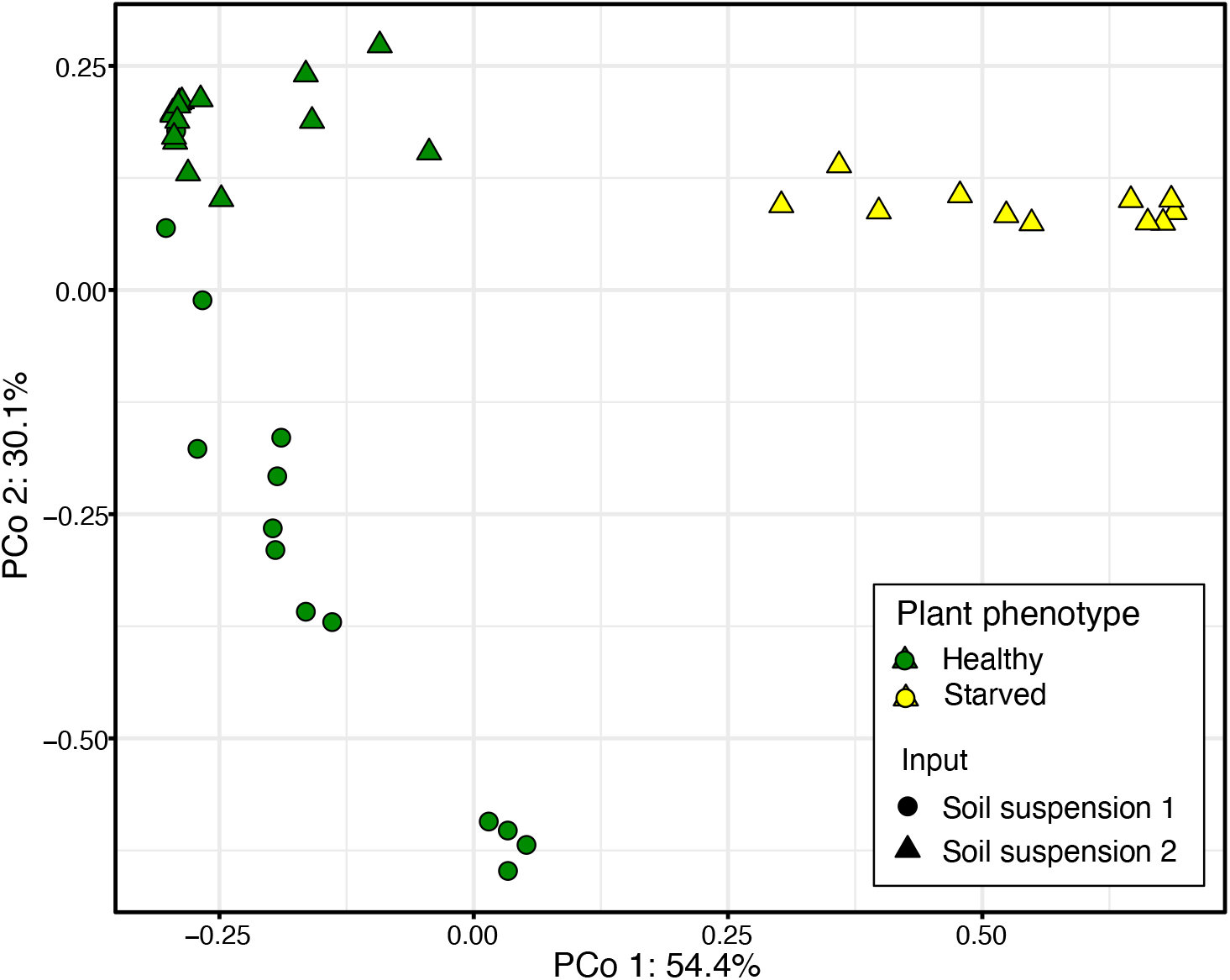
Principal coordinates analysis of *L. burttii* nodules. PCoA plot based on beta diversity calculated using the Bray-Curtis dissimilarity index (32) revealed a clustering of common sample types and a separation of dissimilar sample types.

### Bacterial composition of the nodule microbiome

Both soil suspension types were dominated by Alphaproteobacteria and Gammaproteobacteria. To determine the bacterial composition of the nodule microbiome we estimated the relative abundance at an ASV level. The nodule microbiome of all *Lotus* species was dominated by ASVs belonging to the order Rhizobiales. Nodules of healthy plants were largely dominated by *Mesorhizobium*, independent of the host and soil suspension inoculum. However, while nodules from plants inoculated with soil suspension 1 were colonised with a variety of different *Mesorhizobium* ASVs, the nodules of healthy plants inoculated with soil suspension 2 were almost exclusively colonised by *Mesorhizobium* ASV1 (**Figure 4**). This disparity in *Mesorhizobium* ASV presence is despite the observation that there is no significant difference between the *Mesorhizobium* ASVs present in the two suspensions (*Meso*. Soil S1 v *Meso*. Soil S2, P(>F) = 0.479). Nodules of starved *L. burttii* plants were largely colonised by bacteria belonging to what was taxonomically defined as *Allorhizobium-Neorhizobium-Pararhizobium-Rhizobium* and will be referred to as *Rhizobium* (**Figure 4A**). This suggests that *L. burttii* plants are less selective compared to *L. corniculatus* and *L. japonicus* and develop an ineffective symbiosis with *Rhizobium* strains.

**Figure 4.**
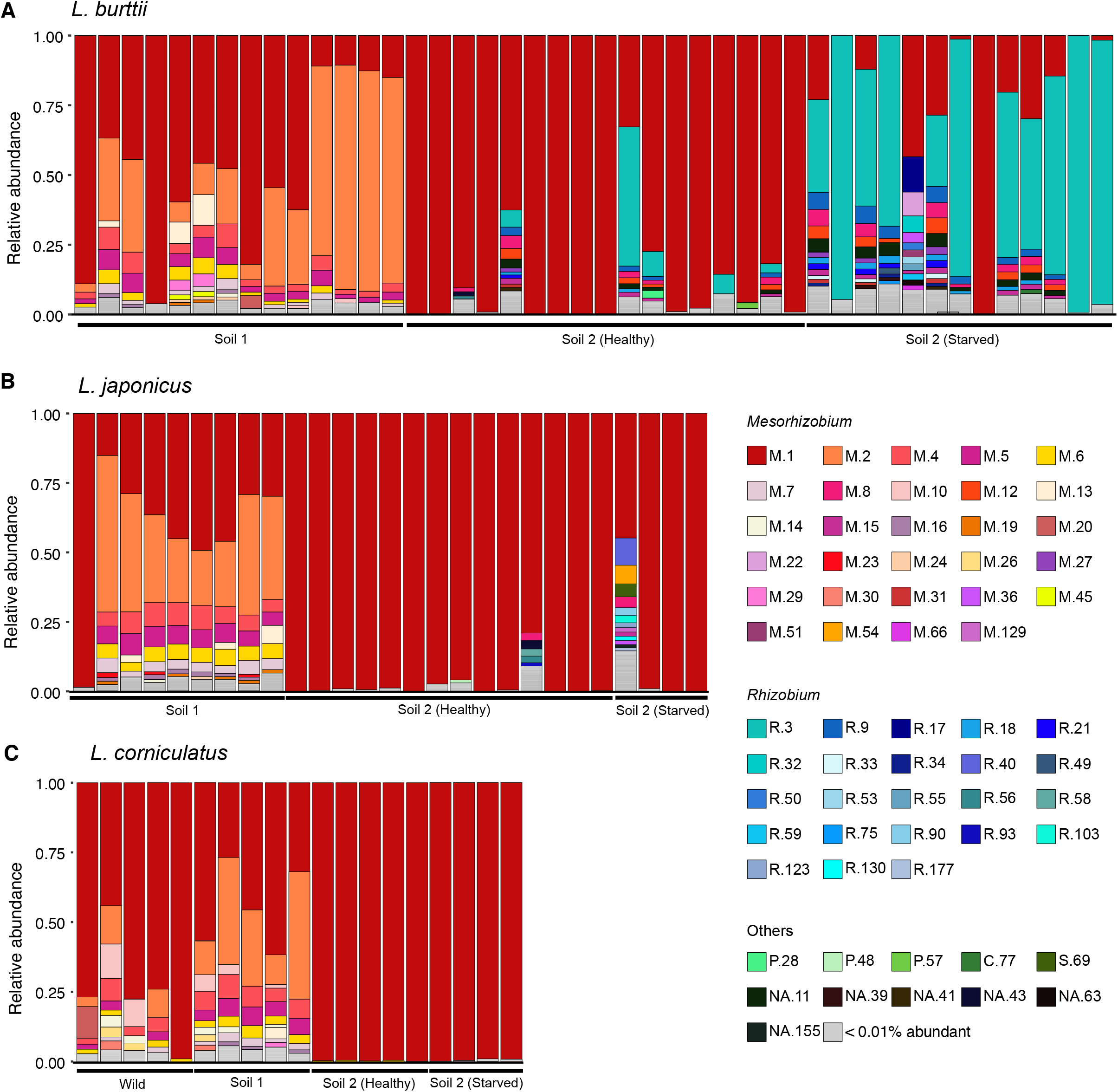
Community profile showing the relative abundance of Amplicon Sequence Variants (ASVs) present in *Lotus* nodules. The relative abundance of ASVs was estimated for *L. burttii* **(A)**, *L. japonicus* **(B)**, and *L. corniculatus* **(C)** using transformed data and the *phyloseq* V 1.26.1 package in R (31). *Mesorhizobium* (M) ASVs are depicted in red, yellow, orange, pink, and purple shades, *Rhizobium* (R) ASVs are depicted in cyan and blue shades. Other ASVs are depicted in green and black. ASVs less than 0.01% abundant are coloured grey. NA, not assigned (taxonomy could only be defined to a Family level).

### *Pseudomonas* are more prevalent in healthy plant nodules

Support Vector Machine (SVM) is a machine learning method used to separate a data set using a linear or non-linear surface (48). In this instance we used a linear-kernel to transform the data and then based on this transformation defined a boundary separating data points, ASVs, based on nodule phenotype of *L. burttii* plants inoculated with soil suspension 2. We used the relative abundance table of ASVs with at least 50 reads from *L. burttii* plants inoculated with soil suspension 2 to construct a linear model using SVM algorithm. 5-fold cross-validation was used to evaluate the model. The model was trained with 70% of the data and evaluated by 30% of the data 5 times (mean of accuracy= 0.89). To identify signature ASVs characteristic of certain sample types, coefficient values for each ASV in the model was obtained using SVM_coef_ function in sckit-learn packages in python (**Figure 5**). The SVM model revealed that *Mesorhizobium* ASV 1 (M.1) was by far the most dominant indicator of a healthy nodule, which is to be expected as *Mesorhizobium* is the typical symbiont of *L. burttii* (49). The second two most influential indicators of a healthy microbiome were *Pseudomonas* ASVs 28 and 57 (P.28 and P.57), which were present in both soil suspension inputs. The 3 ASVs most indicative of a starved *L. burttii* nodule microbiome were *Rhizobium* ASVs.

**Figure 5.**
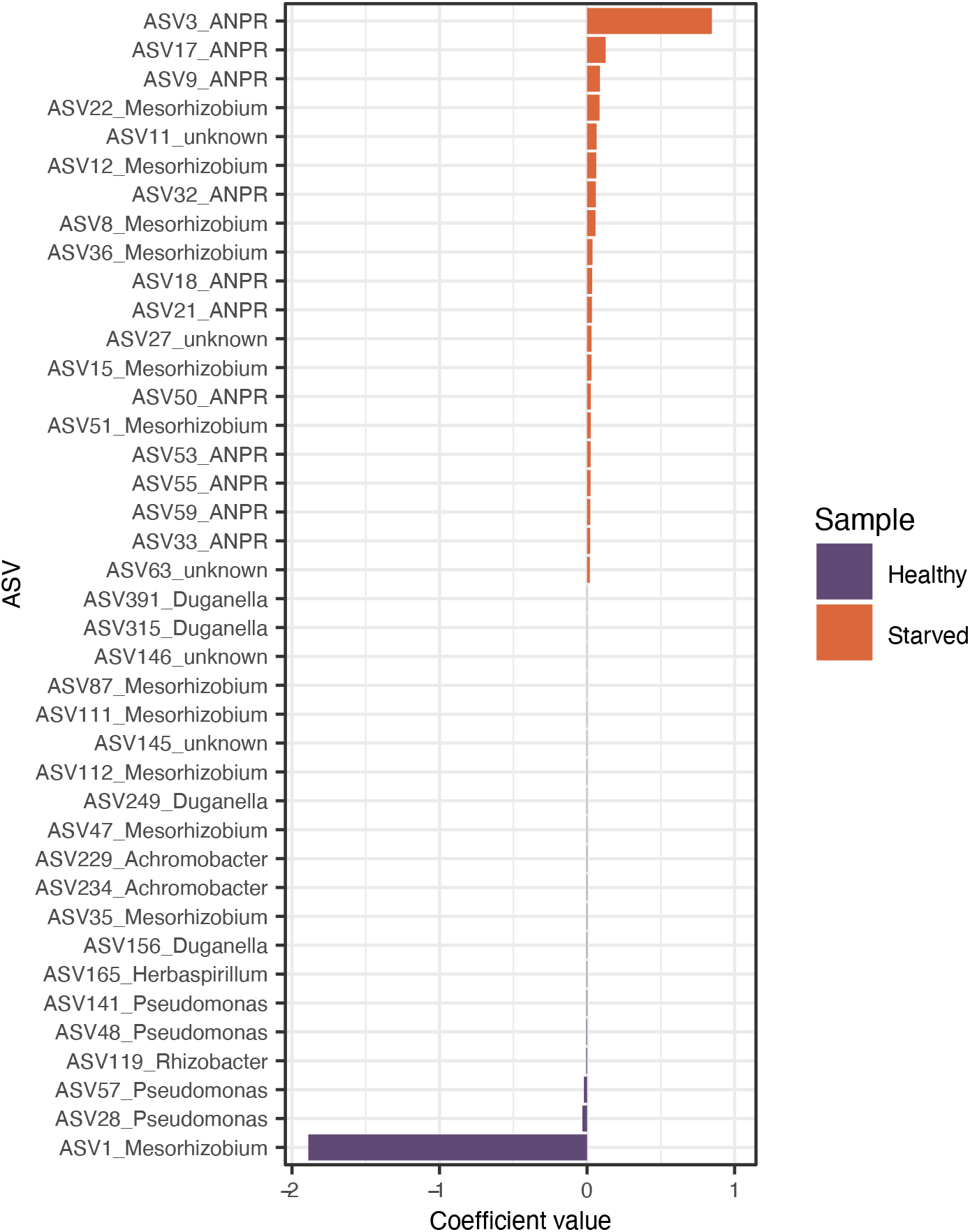
Indicator ASVs of samples. SVM linear model from scikit-learn packages were used to identify separator ASVs between healthy and starved plants. Histogram represents the coefficient scores of top 20 ASVs form heathy and starved plants. Negative coefficient values (purple bars) represent indicator ASVs in healthy plants while positive values (orange bars) show indicator ASVs in starved samples.

### Co-occurrence network of the microbiome

To assess the interaction between and within ASVs from different genera we constructed a microbial network using SparCC (35) algorithm in Fastspar (36) (**Figure S6**). Only significant correlation (|R|>0.2, *P<*0.01) of *Rhizobiaceae* and *Pseudomonadaceae* families are shown in the network (**Figure 6**). The nodes of this network corresponding to ASVs are grouped and coloured by genus. The size of the nodes represents the relative abundance. Each edge between two ASVs represents either positive (orange) or negative (grey) correlations. The ratio of negative to positive interactions within *Rhizobium* ASVs was 1.13 and 1.05 within *Mesorhizobium*, whereas ASVs belonging to *Pseudomonas* are only positively correlated (number of edges=8). The ratio of negative to positive interactions between *Rhizobium* and *Mesorhizobium* (ratio=1.64) was higher compared to this ratio among *Pseudomonas* and *Mesorhizobium* (ratio=0.77). The network illustrated that between *Pseudomonas* and *Rhizobium* ASVs, all correlations were negative (number of edges=37).

**Figure 6.**
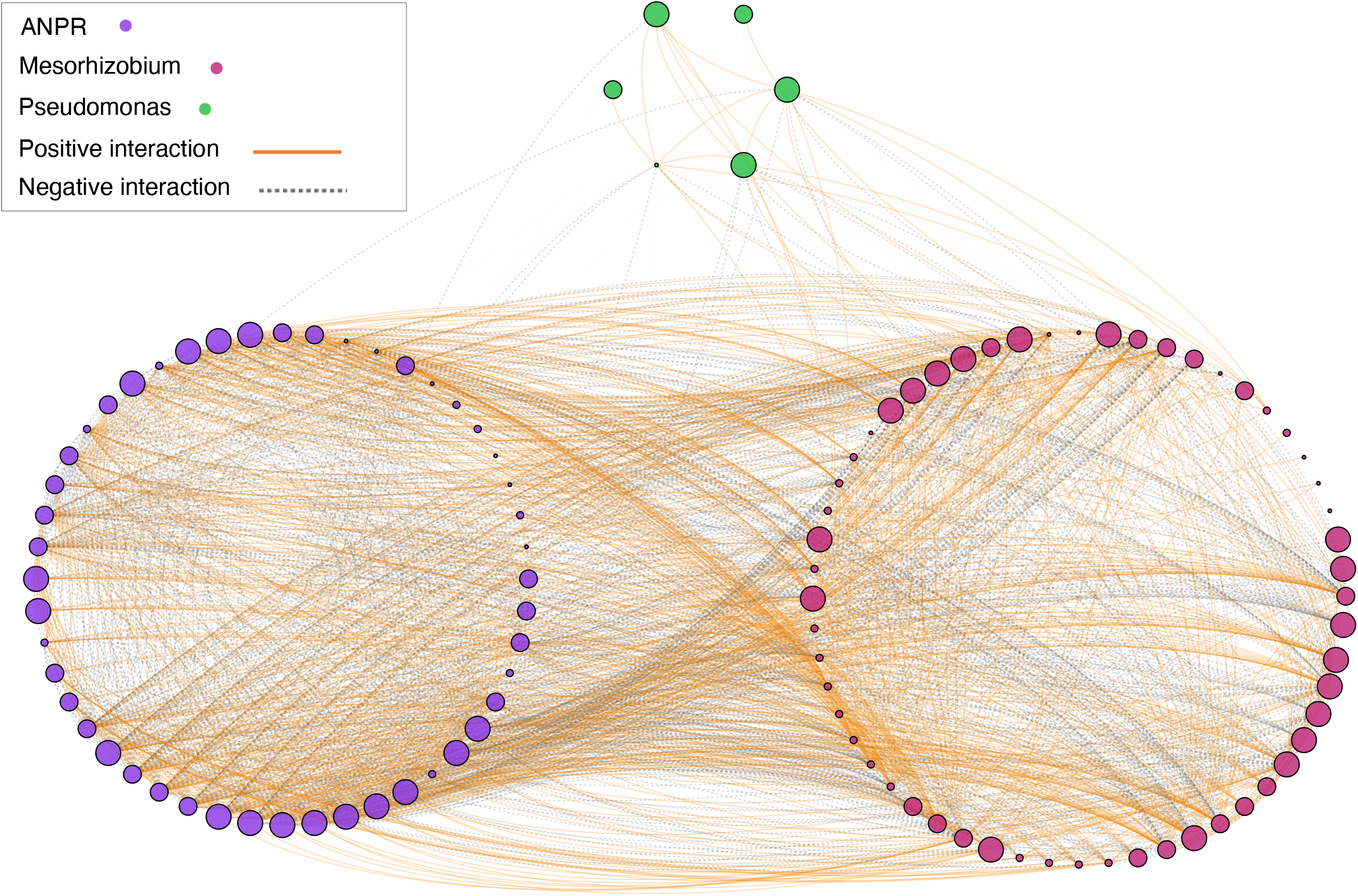
Microbial co-occurrence network of *L. burttii*. An ASV table of *L. burttii* samples was used to infer a correlation network SparCC (35) algorithm implemented in FastSpar (36)tool. The nodes (dots) of this network corresponding to ASVs that are grouped and coloured by Genus. Node size indicates the relative abundance. Each edge (line) between two ASVs represent either a positive (orange line) or negative (grey-dashed line) correlation. Only significant correlations (|R|>0.2, *P <* 0.02) of *Rhizobiaceae* and *Pseudomonadaceae* families are shown in the network.

## DISCUSSION

Nodules of legumes are not only colonized by rhizobia. Despite this, little is known about how microbes other than rhizobia affect the root nodule symbiosis, in particular nodule function and plant health. Here, we characterised variation in the bacterial microbiome of nodules dependent on plant species and soil suspension inoculum and determined correlations between the microbiome makeup and plant health using 16S rRNA gene amplicon sequencing. Our study reveals that (i) the nodule microbiome of *L. japonicus, L. corniculatus*, and *L. burttii* is dependent on soil suspension inoculum, (ii) the nodule microbiome of starved *L. burttii* plants differs to that of the healthy, and (iii) *Pseudomonas* strains are more prevalent in healthy plant nodules then in starved plant nodules.

### Soil suspension inoculum influences *Lotus* spp. nodule microbiome

The nodule microbiome of *Lotus* plants is dependent on the soil suspension inoculum (**Fig. S5**; **Table S3**). Soil is the main influencing factor on the rhizosphere, root, or nodule microbiomes in non-legumes (43) (44) and legumes, such as *Medicago truncatula* (45) and soybean (46, 47). However, many of these studies cite vast differences in the diversity of the microbial communities or physicochemical properties of the soil inputs as the reason for the disparity in plant microbiomes (43, 45, 47). Our results showed that the nodule microbiomes of plants inoculated with different soil suspensions varied significantly (**Table S3**). Also, plants grown in soil suspension 1 produced, on average, more nodules and had a broader range of shoot growth than those inoculated with soil suspension 2 (**Figure S2; Figure 1**). However, the original soil suspensions inoculated onto the plants showed no differences in alpha diversity and only slight although not significant differences in beta diversity (**Figure 2**; **Table S3**). The soils from which the suspensions were produced also had no noteworthy differences in their physicochemical properties (**Table S1**). This suggests that lowly abundant soil microbes, that do not sway diversity measures, may play a pivotal role in how the microbiome functions as a whole. Such a phenomenon has been described in peat soil, where a *Desulfosporosinus* sp., which comprised only 0.006 % of the total microbiome, acted as an important sulphate reducer in the biogeochemical process that diverts carbon flow from methane to CO_2_ (48). Also, *Bacillus* spp., typically found at a low abundance in the rhizosphere compared to *Rhizobia*, increase the number of nodules and/or the size of nodules in legumes (49-52). One striking difference between the nodule microbiomes of *Lotus* plants inoculated with different soils was the number of *Mesorhizobium* ASVs. Many *Mesorhizobium* ASVs were found in plants inoculated soil suspension 1, while those inoculated with soil suspension 2 were almost completely dominated by *Mesorhizobium* ASV M.1 (**Figure 4**). This is despite variation in the beta diversity of *Mesorhizobium* ASVs between the soil inputs was not significant (**Table S3**). Zhang *et al*., showed that the competition between *Mesorhizobium* spp. to nodulate *Cicer arietinum* L. (chickpea) varied dependent on the soil substrate conditions. The native symbiont of chickpea was found to dominate nodule occupation in the native soil, however non-native species dominated in sterilised soils, indicating that other soil microbes contribute to nodulation by the native strain (53). We postulate that lowly abundant taxa may similarly have a disproportionate influence on the overall function of the soil microbiome, in this case impacting nodulation frequency and symbiotic partnerships.

The host genotype had less of an impact on the nodule microbiome than the soil suspension inoculum (**Table S3**). Plant genotype influence on plant-related microbiomes varies among legume species. Brown *et al*., reported that the rhizosphere microbial community of *M. truncatula* is mainly influenced by soil type, while the host genotype is the main determinant of the microbiota in internal plant compartments (45). Similarly, the nodule microbiome of soybean and alfalfa are primarily influenced by host genotype, while the rhizosphere microbiome is more regulated by soil type (52). Contrastingly, the nodule microbiome of cowpea is shaped more by soil type than by plant genotype (54). We found that the influence genotype had on *Lotus* spp. nodule microbiomes was most apparent when also considering plant health phenotype. Healthy *Lotus* spp. plants inoculated with soil suspension 1 had largely similar nodule microbiomes, except when comparing *L. corniculatus* and *L. japonicus* inoculated with soil suspension 1 (**Table S3**). Soil suspension 1 inoculated plants produced exclusively healthy plants, all with nodule microbiomes dominated by *Mesorhizobium* ASVs (**Figure 4**). The significant difference seen between *L. japonicus* and *L. corniculatus*, coupled with the lack of difference seen between them and *L. burttii* indicates there is a variation in nodulation specificity dependent on genotype. We also found that some *Mesorhizobium* ASVs above 0.01% relative abundance were shared exclusively between *L. burttii* and *L. corniculatus* as well as between *L. burttii* and *L. japonicus*. However, no *Mesorhizobium* ASVs were found exclusively in *L. japonicus* and *L. corniculatus*. This suggests that *L. burttii* is the least specific when it comes to establishing root-nodule symbiosis. Host genotype dependent variation in nodulation between these *Lotus* spp. has also been seen upon inoculation with mutant strains that lack the acetylated fucosyl group on the Nod factor of *Mesorhizobium loti*. Nodulation of *L. burttii* was unaffected, while *L. japonicus* and *L. corniculatus* exhibited delayed nodulation and reduced infection (55). Taken together, our results support that the reduced specificity exhibited by *L. burttii* during root-nodule symbiosis allows for a broader range of beneficial *Mesorhizobium* to colonise its nodules.

### Starved *L. burttii* plant nodules harbour a microbiome different to that of healthy plants

*L. burttii* is the only species that we tested that showed a significant difference between the nodule microbiome of heathy and starved plants. Nodules of starved *L. burttii* plants were dominated by *Rhizobium* ASVs, while the nodules of healthy plants were predominantly colonised by *Mesorhizobium* ASVs. *L. burttii* is known to form infected, but ineffective, nodules upon inoculation with *Rhizobium leguminosarum* Norway, however this does not form nodules on *L. japonicus* or *L. corniculatus* (39). This correlates with the observation that starved *L. japonicus* and *L. corniculatus* harboured nodules that were not dominated by *Rhizobium* but rather by *Mesorhizobium* ASVs, similar to the microbiome of healthy plants (**Figure 4**). The variation in starved-plant nodule microbiome between *Lotus* spp. may be explained by how readily each plant is nodulated. Liang *et al*., described that ineffective *R. leguminosarum* Norway colonises nodules of *L. burttii* via cracks in the epidermis, which is not seen in wild type *L. japonicus* (22). This may make *L. burttii* more susceptible to less specific infections and thus increase its vulnerability to forming ineffective symbiosis. This reduced specificity by *L. burttii* is also highlighted in the number of starved plants that contained nodules. 73.8% of starved *L. burttii* plants grew nodules, much more than in *L. japonicus* and *L. corniculatus* (**Figure S3**). The higher frequency of nodulation coupled with the reduced specificity that *L. burttii* exhibits in choosing a nodulation partner leaves the plant susceptible to investing energy in an ineffective nodulation partner, resulting in the starvation of the plant. Conversely, *L. corniculatus* and *L. japonicus* do not exhibit this same level of promiscuity, which is reflected in their nodules being dominated by *Mesorhizobium* in all sample types. The reason as to why a starved plant would harbour a nodule microbiome similar to that of a healthy plant remains to be elucidated. We postulate that it may be simply a delay in the establishment of a successful symbiosis or due to being colonised by non-nitrogen-fixing *Mesorhizobium* strains.

### *Pseudomonas* ASVs are more prevalent in healthy plant nodules

Although the microbiota of all nodule types was dominated by Rhizobiales bacteria, there was a small contingent of non-Rhizobiales ASVs detected as well (**Figure 4**). This is not uncommon in legume nodules, as non-rhizobiales bacteria are often isolated from nodules. Along with Alphaproteobacteria, Betaproteobacteria, Gammaproteobacteria, and Actinobacteria have all been found in various legumes nodules (11, 12, 56-58). Of the non-rhizobia that were present in *Lotus* nodules, *Pseudomonas* was the most prevalent. *Pseudomonas* are often found inside legume nodules (13, 59, 60). We found that *Pseudomonas* ASVs were characteristic of healthy, but not of starved *L. burttii* nodules (**Figure 5**) suggesting they have the potential to support plant health. A *Pseudomonas* strain isolated from *Sophora alopecuroides* promotes plant growth upon reinoculation with *Mesorhizobium* (15). Some *Pseudomonas* spp. can promote leguminous-plant growth via siderophore production, phosphate solubilisation and indoleacetic acid production (12, 15, 16, 61). *Pseudomonas* spp. can indirectly improve plant fitness via antagonistic behaviour towards phytopathogenic fungi (62, 63). Antibiotic compounds production can be induced in *Pseudomonas* through microbe-microbe interactions in soil bacteria (64). We posit that potential microbe-microbe interactions involving *Pseudomonas* also influence the outcome of the root-nodule symbiosis. To analyse any potential microbe-microbe interactions within the nodules we looked for interactions between nodule ASVs. Network analysis comparing the nodule microbiome of healthy and starved *L. burttii* plants revealed significant negative correlations between *Pseudomonas* ASVs and multiple *Rhizobium* ASVs (**Figure 6**). This contrasts with the majority of recorded interactions between *Pseudomonas* and *Rhizobia*, which are synergistic. Interactions can be direct, for example filtrates from *Rhizobium* spp. increasing the cell density of *Pseudomonas fluorescens* (65), or mediated via the plant, for example indoleacetic acid produced by *Pseudomonas* spp. results in a more extensive root system in *Galega officinalis* and an increased number of potential infection sites for the compatible *Rhizobium* sp. (66). The negative correlation we observe between *Pseudomonas* and *Rhizobia* ASVs in *L. burttii* nodules may also be due to an indirect effect mediated by *Mesorhizobium*. Negative correlations were found between *Mesorhizobium* ASVs and *Rhizobia* ASVs. This can be explained by both bacteria competing for nodule colonization. Significant positive correlations were apparent between *Pseudomonas* and *Mesorhizobium* ASV M.1, which is dominant in healthy nodules of plants inoculated with soil suspension 2 (**Figure 4**). Positive interactions occur after the co-inoculation of *Pseudomonas* isolates with a *Mesorhizobium* sp. strain, which leads to an increase in nodule number in chickpea (14). These network interactions coupled with *Pseudomonas* being a known plant-growth-promoting bacterium leads to the hypothesis that these *Pseudomonas* ASVs are beneficial for the plant.

Our results add to the growing assertion that non-rhizobiales bacteria can also contribute to a functional root-nodule symbiosis and healthy plant growth (13). Also, that the nodule microbiome of *Lotus* spp. can vary dependent on soil input, however variations that correlate with plant health are host genotype specific. This research will aid the construction of synthetic communities capable of recreating observed patterns in a bid to narrow down which soil microbes and which microbe-microbe interactions are pivotal in forming the ideal microbiome to maximise plant growth. Refinement of synthetic communities will endeavour towards the development of beneficial plant inocula.

## Supporting information

Figure S1. Reproducibility of plant growth experiments.

Figure S2. Number of nodules per plant after inoculation with soil suspensions.

Figure S3. Nodulation phenotype of starved plants.

Figure S4. Rarefaction curves of sequencing data.

Figure S5. Global PCoA comparison of all samples.

Figure S6. Overview network analysis.

Table S1. Physicochemical analysis of soil samples.

Table S2. Metadata of all samples.

Table S3. PERMANOVA analysis of beta diversity in all nodule microbiome sample types.

## Acknowledgements

This work was supported by the German Research Foundation grant MA 7269/2-1 and Schl446/38-1.

## Competing Interests

The authors declare no competing interests.

## Notes

### Competing Interest Statement

The authors have declared no competing interest.

