## Supplementary material for "The microbiome of *Lotus* nodules varies with plant health in a species-specific manner": Figure S1. Reproducibility of plant growth experiments.

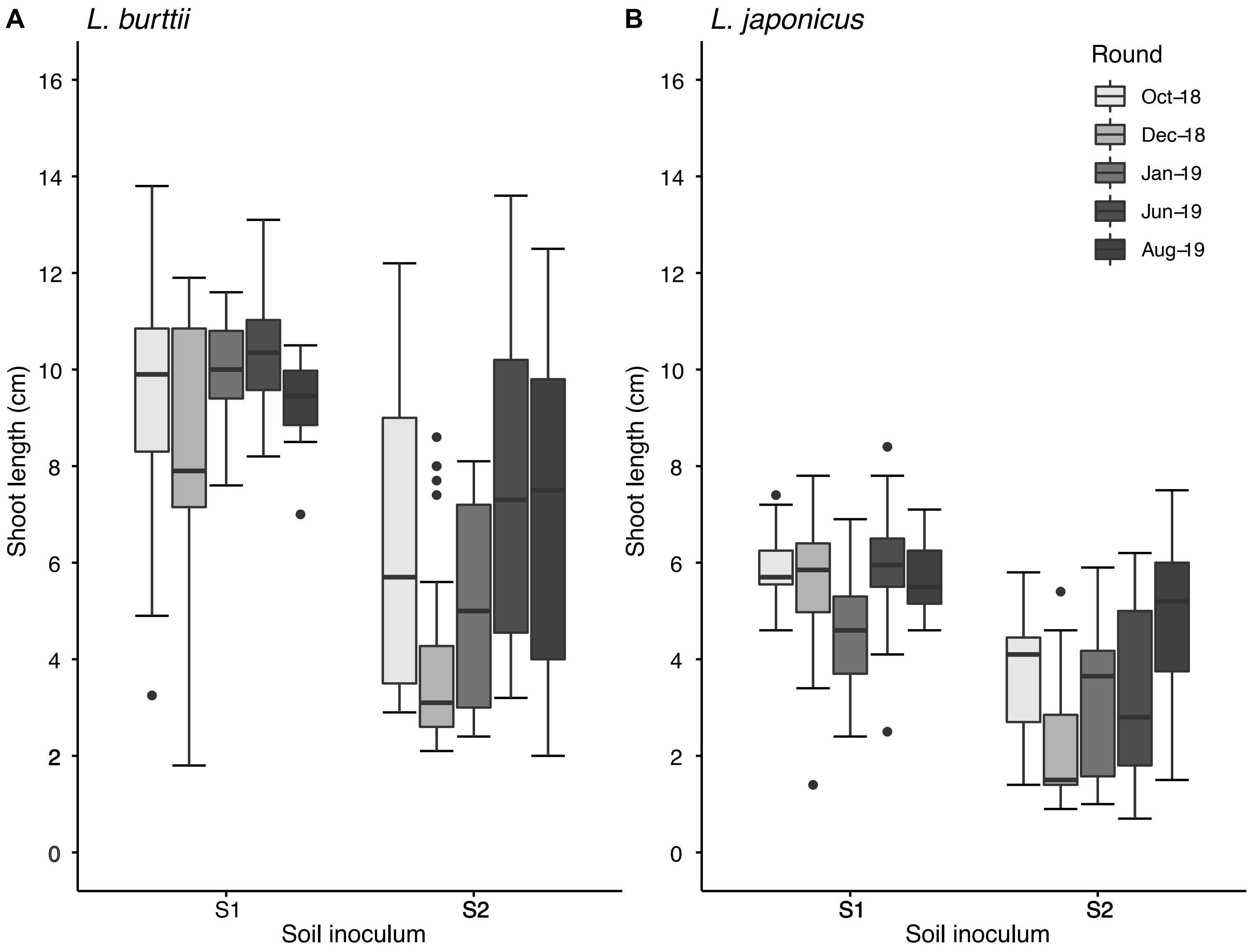


**Figure S1. Reproducibility of plant growth experiments.** Shoot phenotype of **(A)** *L. burttii* and **(B)** *L. japonicus* inoculated with either soil suspension 1 (S1) or soil suspension 2 (S2), grown in closed Weck jars, from independent inoculations. Experiments were carried out in the months indicated with independent soil samples collected in October 2018 and May 2019. Between 50-100 plants were harvested per soil suspension inoculum at 5 weeks post inoculation for each independent experiment.
