## Supplementary material for "The microbiome of *Lotus* nodules varies with plant health in a species-specific manner": Figure S2. Number of nodules per plant after inoculation with soil suspensions.

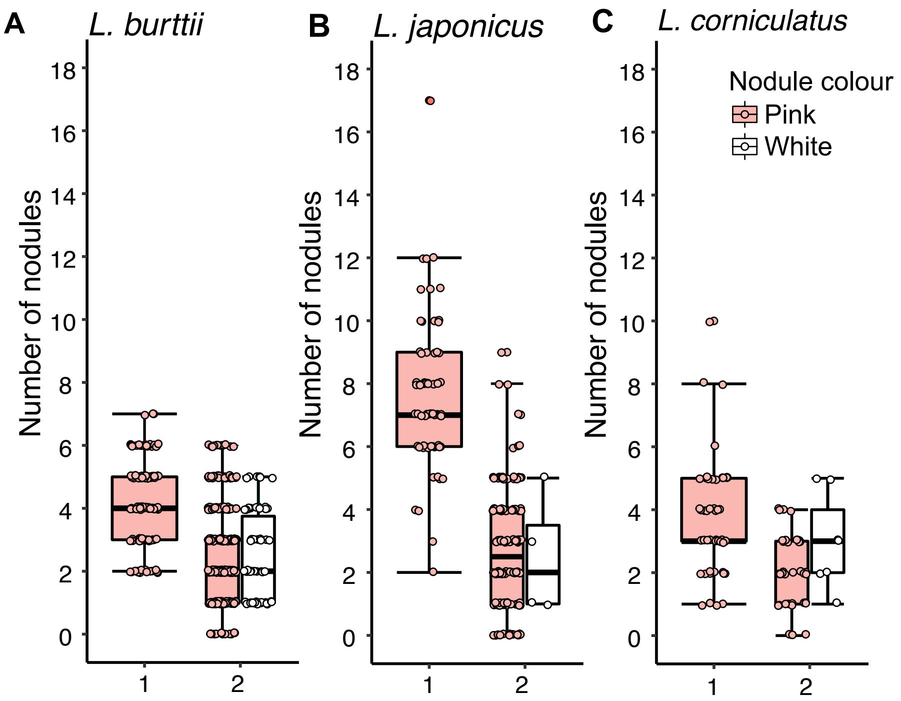


**Figure S2. Number of nodules per plant after inoculation with soil suspensions.** Quantification of pink and white nodules of *L. burttii* **(A)**, *L. japonicus* **(B)**, and *L. corniculatus* **(C)** plants grown in closed Weck jars for 5 weeks after inoculation with soil suspensions 1 (S1) and 2 (S2). Each plot consists of results from two independent experiments. Plants that contained no nodules are not represented. Each sample type contains between 50-150 plants. Mock plants contained no nodules and are not shown.
