## Supplementary material for "The microbiome of *Lotus* nodules varies with plant health in a species-specific manner": Figure S3. Nodulation phenotype of starved plants.

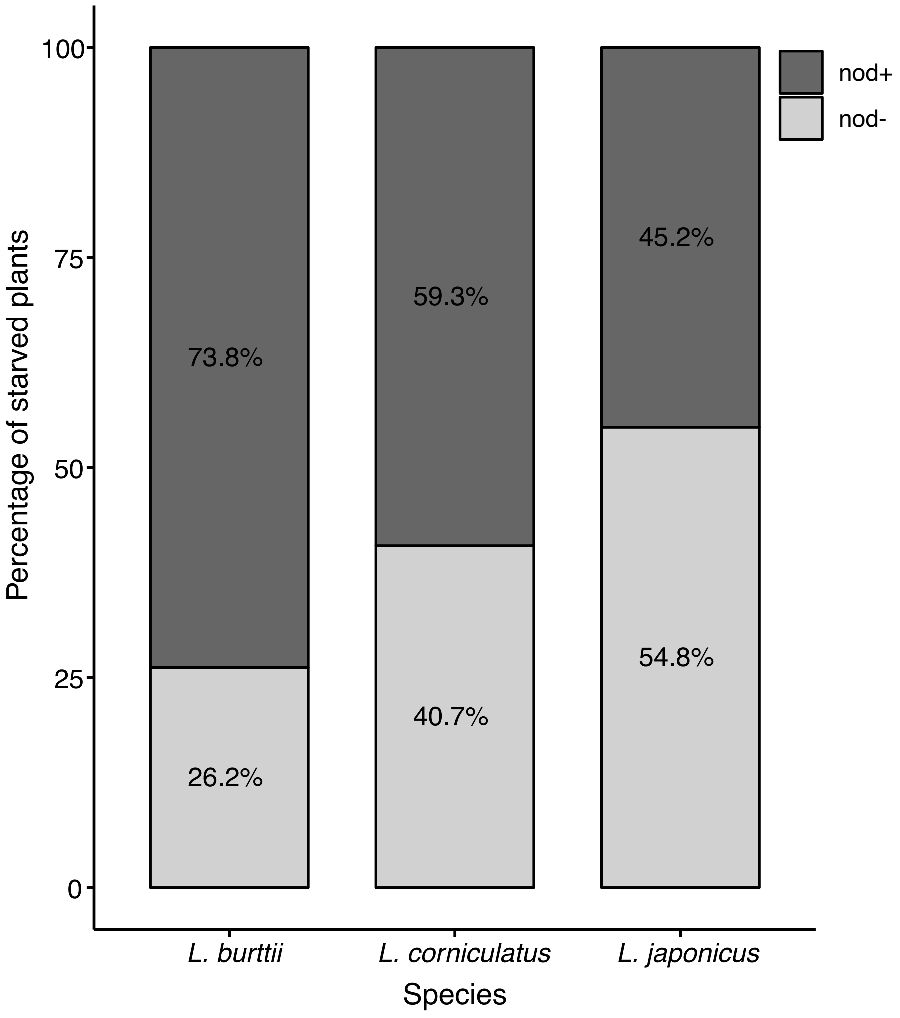


**Figure S3. Nodulation phenotype of starved plants.** Percentages of nodulated plants of *L. burttii* (n=42), *L. corniculatus* (n=27), and *L. japonicus* (n=42) included both pink and white nodules. Plants were inoculated with either soil suspension 1 or 2, grown in closed Weck jars and harvested 5 weeks post inoculation. The plot consists of results from two independent experiments.
