## Supplementary material for "The microbiome of *Lotus* nodules varies with plant health in a species-specific manner": Figure S4. Rarefaction curves of sequencing data.

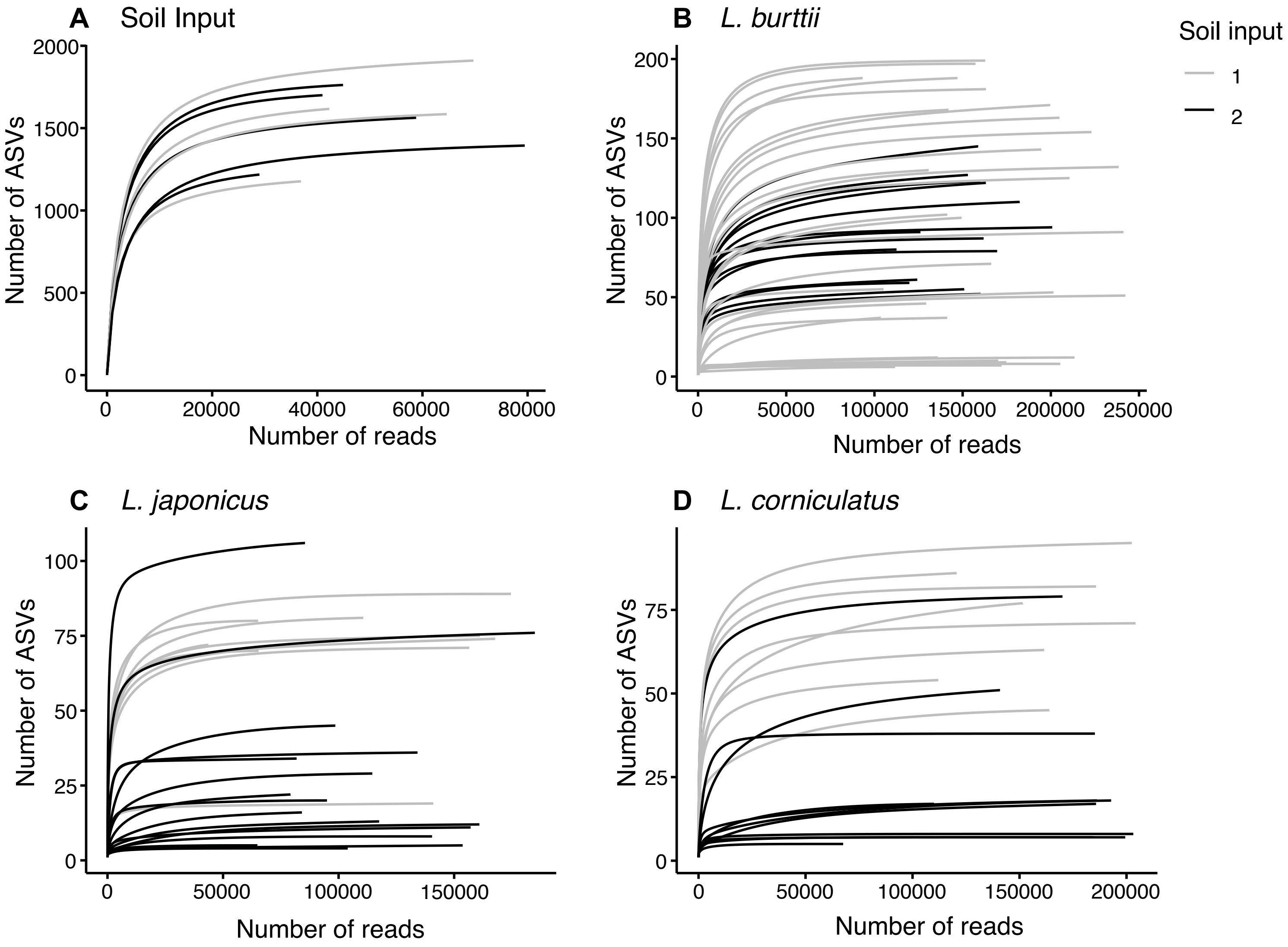


**Figure S4. Rarefaction curves of sequencing data.** Rarefaction curves of nodule samples from each species showing the number of unique ASVs per total reads. Calculated using the *vegan* package in R (34). Soil suspension inoculum for each sample is discern by colour.
