## Supplementary material for "The microbiome of *Lotus* nodules varies with plant health in a species-specific manner": Figure S5. Global PCoA comparison of all samples.

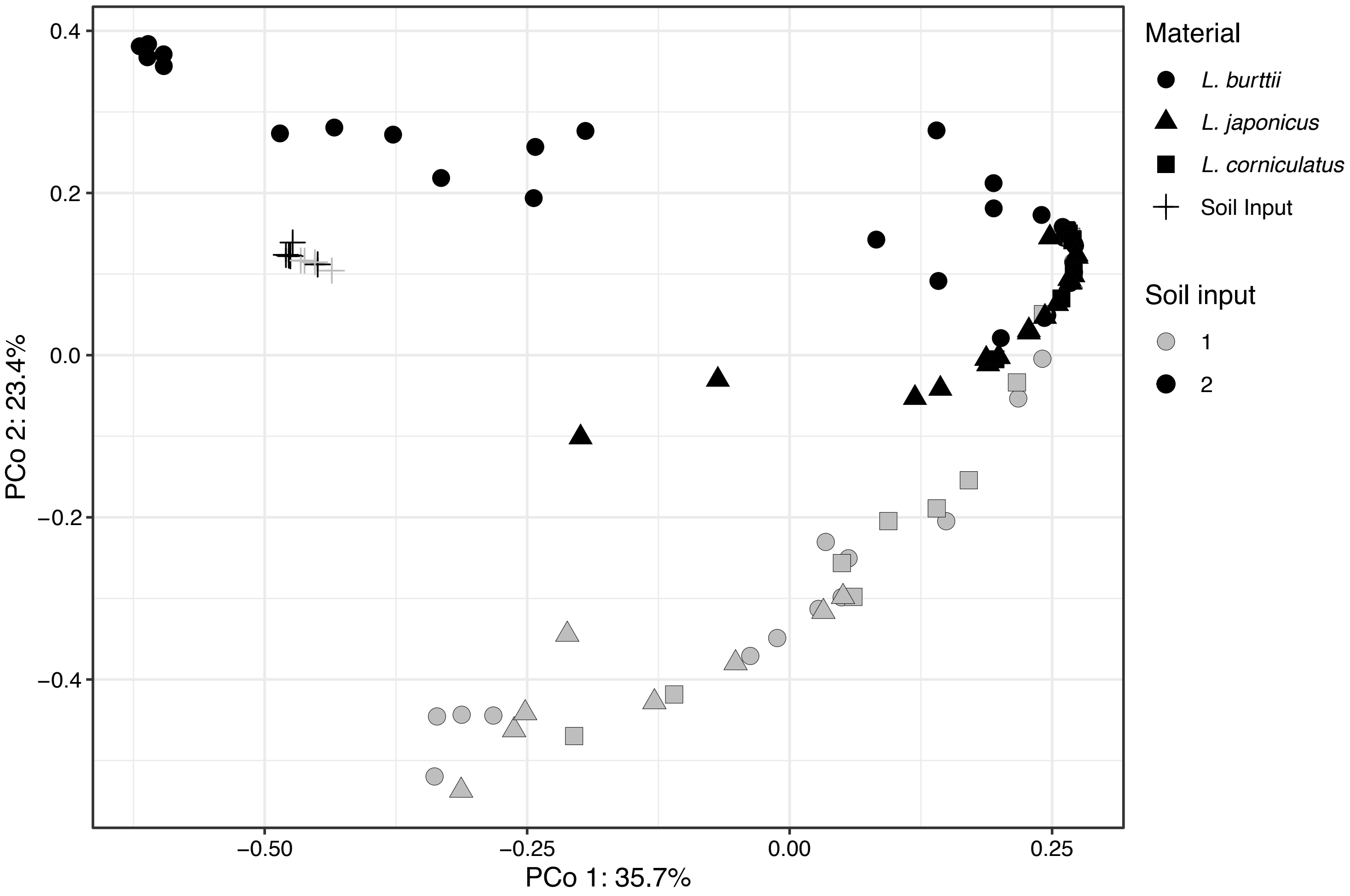


**Figure S5. Global PCoA comparison of all samples.** PCoA plot of all soil suspension input and nodule samples based on beta diversity calculated using the Bray-Curtis dissimilarity index (32).
