## Supplementary material for "The microbiome of *Lotus* nodules varies with plant health in a species-specific manner": Figure S6. Overview network analysis.

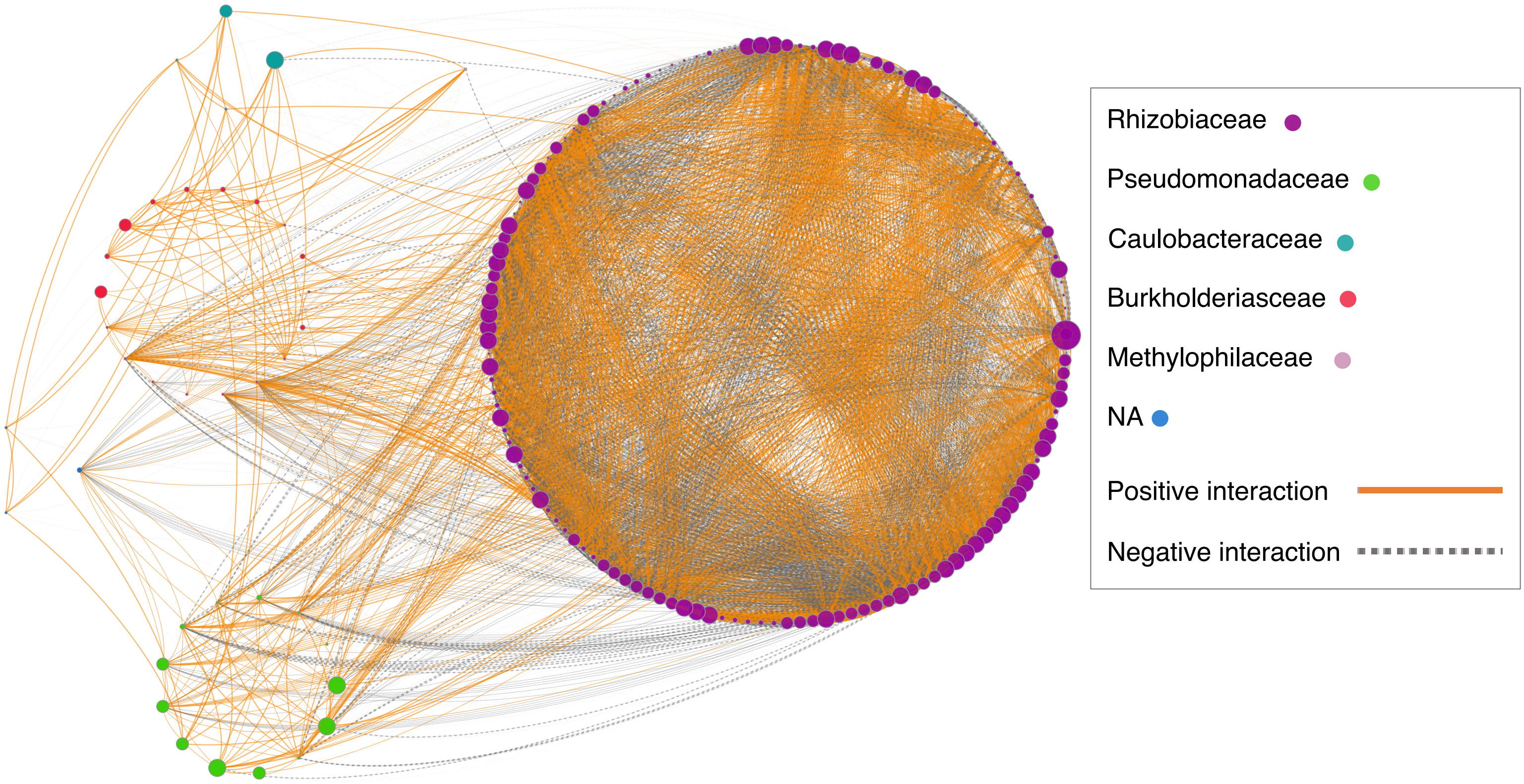


**Figure S6. Overview network analysis**. An ASV table of *L. burttii* samples was used to infer a correlation network SparCC (35) algorithm implemented in FastSpar (36)tool. The nodes (dots) of this network corresponding to ASVs that are grouped and coloured by Family. Node's size indicates the relative abundance. Each edge (line) between two ASVs represent either a positive (orange line) or negative (grey-dashed line) correlation. Significant correlations (|R|>0.2, P < 0.01) are shown in the network.
